# Orchid seed germination through auto-activation of mycorrhizal symbiosis signaling regulated by gibberellin

**DOI:** 10.1101/2023.04.26.538379

**Authors:** Chihiro Miura, Yuki Furui, Tatsuki Yamamoto, Yuri Kanno, Masaya Honjo, Katsushi Yamaguchi, Kenji Suetsugu, Takahiro Yagame, Mitsunori Seo, Shuji Shigenobu, Masahide Yamato, Hironori Kaminaka

**Affiliations:** Faculty of Agriculture, Tottori University, Tottori, Japan; Graduate School of Agriculture, Tottori University, Tottori, Japan; RIKEN Center for Sustainable Resource Science, Yokohama, Japan; Functional Genomics Facility, NIBB Core Research Facilities, National Institute for Basic Biology, Okazaki, Japan; Department of Biology, Graduate School of Science, Kobe University, Kobe, Japan; Mizuho Town Museum, Mizuho, Japan; Faculty of Education, Chiba University, Chiba, Japan; Unused Bioresource Utilization Center, Tottori University, Tottori, Japan

**Author notes:** Corresponding Author Hironori Kaminaka.

**Keywords:** Co-option, Gibberellins, Mycoheterotrophy, Mycorrhizal symbiosis, Orchidaceae, Seed germination

## Abstract

Orchids parasitically depend on external nutrients from mycorrhizal fungi for seed germination. Previous findings suggest that orchids utilize a genetic system of mutualistic arbuscular mycorrhizal (AM) symbiosis to establish parasitic symbiosis. In AM symbiosis, recent studies have revealed that the plant hormone gibberellin (GA) negatively affects fungal colonization and development. Although previous studies imply that GA is important for orchid mycorrhizal symbiosis, the molecular mechanism of seed germination in which mycorrhizal symbiosis co-occurs remains unclear because, in AM plants, GA regulates seed germination and symbiosis positively and negatively, respectively. To elucidate this conflict, we investigated the effect of GA on *Bletilla striata* seed germination and mycorrhizal symbiosis using asymbiotic and symbiotic germination methods. Additionally, we compared the transcriptome profiles between asymbiotically and symbiotically germinated seeds. Exogenous GA negatively affected seed germination and fungal colonization, and endogenous bioactive GA was actively converted to the inactive form during seed germination. Transcriptome analysis showed that *B. striata* shared many of the induced genes between asymbiotically and symbiotically germinated seeds, including GA metabolism- and signaling-related genes and AM-specific marker homologs. Our study suggests that orchids have evolved in a manner that they do not use bioactive GA as a positive regulator of seed germination and instead, auto-activate the mycorrhizal symbiosis pathway through GA inactivation to accept the fungal partner immediately during seed germination.

## Introduction

Seed germination is an important process in the plant life cycle because it determines subsequent plant survival and reproductive success (Rajjou et al., 2012). Diverse environmental factors to which seeds are exposed, such as light, temperature, water, oxygen, and chemical substances, break seed dormancy, and stimulate seed germination (Shu et al., 2016). It is well known that the plant hormone gibberellin (GA) plays an essential role in promoting seed germination in many plant species, including model plants, *Arabidopsis*, and many crop plants (Tuan et al., 2018). During seed imbibition of cereal crops, the embryo synthesizes GA, which is subsequently released to aleurone cells to activate the synthesis and secretion of hydrolases such as α-amylase (Tuan et al., 2018). These hydrolases degrade starch and other nutrients stored in the endosperm into small molecules that are used by embryos for supporting seed germination and seedling growth (Tuan et al., 2018).

Orchids, which belong to the family Orchidaceae, produce a large number of small seeds containing 200 µm-long embryos on average with starch-deficient endosperm, which is termed dust-like seeds (Arditti and Ghani, 2000). The seeds consisting only of an undifferentiated embryo surrounded by a thin seed coat, parasitically depend on nutrients supplied by orchid mycorrhizal (OM) fungi for seed germination and seedling development (Eriksson and Kainulainen, 2011). Under natural conditions, the orchid seeds rely on OM associations to obtain carbon, nitrogen, and phosphorus during the early developmental stage, which is called ’initial mycoheterotrophy’ (Leake, 1994; Merckx, 2013). Because orchid seed germination is unique and complex in terms of its requirements (Arditti, 1967; Rasmussen et al., 2015), it is assumed that orchid germination is regulated by a different mechanism from that underlying the germination of the majority of photosynthetic plants. In the context of GA, which is a well-known seed germination stimulator, Van Waes and Debergh (1986) examined the effects on the asymbiotic (aseptic) germination *in vitro* of four western European terrestrial orchids (*Cypripedium calceolus*, *Dactylorhiza maculata*, *Epipactis helleborine*, and *Listera ovata*), showing that GA negatively affected the germination of *D. maculata* and *L. ovata* and had no significant effect on *C. calceolus* and *E. helleborine* (Van Waes and Debergh, 1986). Similarly, GA exerted no stimulatory effect on the asymbiotic seed germination of *Calanthe discolor*, a terrestrial orchid that grows in eastern Asia (Miyoshi and Mii, 1995). Chen *et al* (2020) showed no significant effects of GA on asymbiotically germinated seeds of the epiphytic orchid *Dendrobium officinale*, and inhibition of fungal colonization in symbiotically germinated seeds in the case of applying high concentrations of exogenously GA (Chen et al., 2020). Another report showed the positive effects of GA on asymbiotically germinated immature seeds of *Pseudorchis albida*, probably due to the differences in the degree of seed maturity (Pierce and Cerabolini, 2011). Although the lack of endosperm in the common ancestor of orchids may explain low sensitivities to GA in phylogenetically distant orchids, the molecular mechanisms of seed germination remain unclear.

Orchid mycorrhizal fungi penetrate seeds either through the suspensors (Peterson and Currah, 1990; Richardson et al., 1992; Rasmussen and Rasmussen, 2009) or through rhizoids (Williamson and Hadley, 1970) and then form dense coils of mycelium called pelotons. Orchids obtain nutrients through both live and degenerating pelotons (Kuga et al., 2014). Most orchids, including members of the earliest-diverging clade of the Orchidaceae, form OM symbioses with free-living, saprophytic fungi belonging to the phylum Basidiomycota (Rasmussen, 2002; Yukawa et al., 2009). The Orchidaceae is sister to the other members of the Asparagales, which only form symbioses with the Glomeromycotina (except for orchids) that constitute the most ancient mutualistic plant-fungi association, termed arbuscular mycorrhizae (AM) (Delaux et al., 2013). It is generally accepted that the fungal partner of orchids has shifted from Glomeromycotina to Basidiomycota (Yukawa et al., 2009). Supporting this idea, our previous molecular-based study of *Bletilla striata* has shown that orchids share the molecular components such as common symbiotic genes and their concomitant expression common to AM plants for mycoheterotrophic symbiosis, which implies that orchids utilize a genetic system of AM symbiosis (Miura et al., 2018). Consistent with this suggestion, the common symbiotic pathway genes, involved in a putative signal transduction pathway shared by AM and the rhizobium–legume symbiosis, such as symbiosis receptor-like kinase *SymRK*, calcium- and calmodulin-dependent protein kinase *CCaMK*, and calcium signal decoding protein *CYCLOPS*, are present in other orchid species, *Apostasia shenzhenica*, *Dendrobium catenatum*, *Phalaenopsis equestris*, and *Gastrodia elata* (Radhakrishnan et al., 2020; Xu et al., 2021). In terms of AM symbiosis, recent studies have revealed that abnormal elevation of GA signaling decreases the colonization of the host root by AM fungi (Floss et al., 2013; Foo et al., 2013), and GA reduces hyphal colonization and arbuscule formation during AM symbiosis (Takeda et al., 2015). The arbuscule formation required the presence of DELLA (aspartic acid–glutamic acid–leucine–leucine–alanine) proteins, which are negative regulators of GA signaling (Floss et al., 2013). In *Lotus japonicus*, a complex comprising CCaMK, CYCLOPS, and DELLA binds to the promoter of *REDUCED ARBUSCULAR MYCORRHIZA1*, a central regulator of arbuscule development (Pimprikar et al., 2016). Thus, GA signaling negatively affects AM fungal colonization and development.

More than 80% of the land plant species germinate using nutrient sources stored in their seeds and establish AM symbiosis after root development. This means that the mechanisms of seed germination and mycorrhizal symbiosis are independent of most photosynthetic plants. In contrast, these two events, for both of which GA is a key regulator, co-occur in orchids. Previous orchid transcriptomic studies have reported high expression of genes related to GA biosynthesis, catabolism, and signaling during symbiotic germination (Zhao et al., 2014; Liu et al., 2015; Miura et al., 2018). Given the low sensitivities to GA and the expression patterns of GA metabolic and signaling genes in orchid species, we hypothesized that orchids have evolved to not use GA as a positive regulator of seed germination, in order to establish and maintain symbiotic associations during germination. To elucidate the connecting mechanisms of orchid seed germination and mycorrhizal symbiosis at the molecular level, we investigated the effect of GA on seed germination and mycorrhizal symbiosis mainly by using a terrestrial orchid, *Bletilla striata* cv. *Murasakishikibu* (subfamily Epidendroideae, tribe Arethuseae), which is an initially mycoheterotrophic plant species mostly distributed in China and

Japan, and is usually found in moist grasslands. Because *B. striata* can grow almost synchronously *in vitro* both asymbiotically and symbiotically (Yamamoto et al., 2017), the germination rates and symbiotic levels were examined in the growth conditions with or without GA. As a result, GA negatively affected both seed germination and fungal colonization. The GA-treated asymbiotic germination assay confirmed the negative effects on some phylogenetically distant orchids. Asymbiotically- and symbiotically germinated *B. striata* protocorms were subjected to transcriptome analysis using RNA-sequencing (RNA-seq) to determine whether *B. striata* share differentially expressed genes (DEGs) between the two states. The results showed that *B. striata* shared more than half of the induced DEGs. The commonly upregulated DEGs included AM-specific marker homologs, meaning that symbiosis signaling is activated during seed germination even without fungi. Our study, therefore, concluded that orchids auto-activate the mycorrhizal symbiosis pathway during seed germination through GA inactivation, suggesting an adaptive mechanism to reconcile the two events, seed germination and mycorrhizal symbiosis.

## Materials and Methods

### Plant materials and fungal strain

Seeds of *Bletilla striata* cv. Murasakishikibu and its symbiotic fungus, *Tulasnella* sp. strain HR1-1, were used in this work. The origins of the plant and fungal line have been described in detail previously (Yamamoto et al., 2017). *Spiranthes australis* and *Cremastra appendiculata* var. *variabilis* were collected from Houki town, Tottori Prefecture, Japan, and Kisakata town, Akita Prefecture, Japan, respectively. *B. striata*, *S. australis*, and *C. appendiculata* were grown in a greenhouse, and matured seeds were harvested at five, one, or four months, respectively, after self-pollination. Matured *Goodyera crassifolia* and *Vanda falcata* var. *Beniohgi* seeds were collected from Kami city, Kochi Prefecture, Japan, and Kihoku town, Mie Prefecture, Japan, respectively. Collected seeds were stored at 4°C until they were used. Fungal colonies were cultivated on potato dextrose agar (PDA; Kyokuto, Tokyo, Japan) medium at 25°C until symbiotic germination experiments were required.

### Asymbiotic and symbiotic germination on agar medium

Asymbiotic and symbiotic germination procedures were performed according to the method of Yamamoto et al. (2017), either with or without additional growth regulators: gibberellic acid 3 (GA3) (Nakalai Tesque, Kyoto, Japan), uniconazole-P (FUJIFILM Wako Pure Chemical, Kyoto, Japan), and abscisic acid (ABA) (Sigma-Aldrich, St. Louis, MO, USA) dissolved in ethanol. The method is briefly described below. The seeds were surface sterilized in 1% (v/v) sodium hypochlorite solution for 2 min for *B. striata* and *V. falcata*, 30 s for *S. australis*, 4 min for *C. appendivulata*, and 5 min for *G*. *crassifolia.* After rinsing with sterilized distilled water, 40–800 sterilized seeds derived from one mature fruit capsule of each orchid species per plate were sown onto HYPONeX-sucrose agar (HA) medium (3.0 g Hyponex [6.5–6-19] [Hyponex Japan, Osaka, Japan], 2.0 g peptone, 30 g sucrose, 10 g agar, 1 l distilled water, pH 5.5) for asymbiotic germination or oatmeal agar (OMA) medium (2.5 g Oatmeal agar (Difco, Franklin, New Jersey, USA), 6.5 g agar, 1 l distilled water, pH 5.5) pre-inoculated with a culture of *Tulasnella* sp. for symbiotic germination. Growth regulators were added to the autoclaved media after cooling to 60 °C. Ethanol (0.01%, v/v) was included in the media as a control treatment. The germination experiments were conducted at 25°C in the dark. Germinated seeds were counted or harvested every week for 3 weeks and stored in a formalin–acetic acid–alcohol (FAA) solution at 4 °C for cell staining or at –80 °C after freezing in liquid nitrogen for RNA extraction and phytohormone quantification.

### Quantitative measurement of seed germination and symbiotic cells

The germination was defined as the emergence of a shoot in *B. striata* and *V. falcata* or rhizoids in *S. australis*, *G*. *crassifolia*, and *C. appendiculata*. To measure the germination rate, at least three plates with approximately 100 seeds each, were observed under a stereomicroscope (Olympus SZX16, Tokyo, Japan) in an individual experiment. The measurements were repeated at least two times for asymbiotic and symbiotic germination of *B. striata*, *S. australis*, *G*. *crassifolia,* and *V. falcata*. The experiments for *C. appendiculata* seed germination were performed only twice due to the small number of seeds available in this study. The FAA-fixed protocorms were rinsed with distilled water and softened in 10% KOH at 105 °C for 5 min. The alkalized samples were rinsed with distilled water, neutralized by 2% (v/v) HCl for 5 min, and stained with 10% (v/v) ink dye solution (10% Pelikan 4001 Brilliant Black and 3% acetic acid) at 95 °C for 30 min for counting of symbiotic cells or with 10 µg/ml WGA-Alexa Fluor 488 solution (Thermo Fisher Scientific, Waltham, MA, USA) overnight and 1 µg/ml Calcofluor White solution (Sigma-Aldrich, St Louis, MO, USA) for 15 min in the dark at room temperature for observation of the fungal distribution. Ink-stained materials were soaked in lactic acid (Nacalai tesque, Kyoto, Japan) at 4 °C before microscopic observation. Procedures of symbiotic cell counting were performed according to the method of Yamamoto et al. (2017). The method is briefly described below. The seed coat removed protocorms were held between a glass slide and a cover glass and were lightly crushed. The number of symbiotic cells was counted under a BX53 light microscope (Olympus) and averaged from randomly selected ten individual protocorms. The cell counting experiment was independently repeated four times.

### Asymbiotic germination on filter paper

Sterilized *B. striata* seeds were placed on a filter paper (ø 70 mm) containing 2 ml of a test solution. The seeds of *O. sativa* cv. Nipponbare and *L. japonicus* MG-20 were also seeded on a filter paper using the same procedure. These seeds were incubated at 25°C for three weeks, three days, and two days in the dark for *B. striata*, *Oryza sativa* and *L. japonicus*, respectively. Germination was defined as the emergence of a root in *O. sativa* and *L. japonicus*. The experiment was independently repeated two times, each containing 3–5 replicate plates.

### RNA preparation and gene expression analysis by qRT-PCR

Total RNA was extracted from approximately 5 mg of *B. striata* seeds or protocorms for each condition using the Total RNA Extraction Kit Mini for Plants (RBC Bioscience, New Taipei, Taiwan), following the manufacturer’s protocol. First-strand cDNA synthesis was performed using the ReverTra Ace qPCR RT Master Mix with gDNA Remover (Toyobo, Osaka, Japan) following the manufacturer’s protocol. Quantitative RT-PCR assays were carried out using the THUNDERBIRD SYBR qPCR Mix (Toyobo) on a CFX connect real-time detection system (Bio-Rad Laboratories, Hercules, CA, USA) using the following program: 95°C for 10 min; 45 cycles of 95°C for 30 s, 60°C for 30 s, and 72°C for 30 s; and a final extension at 72°C for 5 min. Fold changes were calculated using the expression of a housekeeping gene, *UBIQUITIN5*, in *B. striata* as the internal control. The gene-specific primer sequences used in this study are shown in Table **S1** and Miura et al. (2018). At least five biological replicates containing three technical replicates each were performed. After the threshold cycle (*Ct*) values were averaged, the fold changes were calculated using the delta-delta cycle threshold method (Livak and Schmittgen, 2001) or the relative standard curve method (Pfaffl, 2001).

### Quantification of endogenous GA levels in *B. striata*

Asymbiotically- or symbiotically germinated *B. striata* protocorms were harvested two weeks after seeding and stored at –80 °C after freezing in liquid nitrogen. The seeds, before seeding, were also stored at –80 °C as a control sample. The frozen samples were weighed after lyophilization. After grounding and homogenizing, the 35.15–234.89 mg samples were subjected to LC-MS/MS to quantify the endogenous GAs according to the method of Kanno et al. (2016). Data were obtained from three independent replicate experiments.

### RNA-sequencing and data analysis

The total RNA of asymbiotically germinated protocorms (AP) was treated with RNase-free DNase I to remove residual genomic DNA and cleaned using the RNeasy Mini Kit (Qiagen, Hilden, Germany) according to the manufacturer’s protocol. The quality and quantity of the purified RNA were confirmed by measuring its absorbance at 260 nm and 280 nm (A260/A280) using a NanoDrop ND-1000 spectrophotometer (Thermo Fisher Scientific, Waltham, MA, USA) and by electrophoresis using an Agilent 2100 Bioanalyzer (Agilent Technologies, Santa Clara, CA, USA). RNA-seq library was constructed from the total extracted RNA using the Illumina TruSeq RNA Library Prep Kit (Illumina, San Diego, CA, USA) according to the manufacturer’s protocol. Multiple cDNA libraries were sequenced using the Illumina HiSeq platform with 100 bp or 125 bp single-end reads. Three biological replicates were prepared for the transcriptome analysis. Consequently, more than seven million raw reads per sample were obtained (Table S2). Low-quality reads (< Q30) and adapter sequences were filtered and trimmed using Fastp (Chen et al., 2018).

### Differential expression analysis

For data analysis, raw reads of asymbiotically geminated *B. striata* generated in this study (the DNA Data Bank of Japan (DDBJ) DRA accession DRR439921–DRR439932) and symbiotically germinated *B. striata* obtained from our previous study (DDBJ DRA accession DRR099058–DRR099075) (Miura et al., 2018) were used. The reads were mapped using Bowtie 2 (Langmead and Salzberg, 2012) and the transcript abundance was estimated using eXpress ver. 1.5.1 (Roberts and Pachter, 2012) in accordance with the method of Miura et al. (2018). Differences in library size were corrected using the trimmed mean of M-value normalization method, and EdgeR (Robinson et al., 2010) was used to identify DEGs with a Log2 fold change (FC) of ≥ 1.0 or ≤ −1.0 and false discovery rates (FDR) <0.05. The time-course data were analyzed using a general linear model. To identify the DEGs in germinated and AM-colonized rice, publicly available short-read data were obtained from the Short-Read Archives at the National Center for Biotechnology Information BioProject accession PRJNA474721 (Narsai et al., 2017) and DDBJ BioProject accession PRJDB4933 (Kobae et al., 2018), respectively. The downloaded raw reads were filtered and trimmed using Fastp with the option “-q 30” (Chen et al., 2018), and then mapped using STAR (Dobin et al., 2013). The reference genome (Oryza_sativa IRGSP-1.0.48) was downloaded from Ensembl plants (https://plants.ensembl.org/index.html). The aligned reads were counted using featureCounts (Liao et al., 2014); subsequently, the genes were subjected to differential expression analysis using the EdgeR package (Robinson et al., 2010). Differentially expressed genes were defined as Log2 FC ≥ 1.0 or ≤ –1.0 and FDR < 0.05.

### Gene annotation and gene-ontology analysis

The *B. striata de novo* reference assembly (Miura et al., 2018) was functionally annotated with EnTAP (Hart et al., 2020) using the NCBI NR, Plant RefSeq, and UniProt (https://www.uniprot.org/) databases. The gene-ontology (GO) enrichment analysis of DEGs was performed using the topGO package in the R environment with Fisher’s exact test (FDR < 0.05).

## Results

### Gibberellin negatively affects both fungal colonization and seed germination in *B. striata*

Surface-sterilized *B. striata* seeds were sown on an OMA medium containing 1 µM GA inoculated with the previously isolated *Tulasnella* sp. HR1-1 (hereafter *Tulasnella*) (Yamamoto et al., 2017) to examine the effects of GA on symbiosis. No significant effect was found on the growth of GA-treated *Tulasnella* (Supplemental Figure S1A). Since a protocorm colonized with *Tulasnella* contains dozens of symbiotic cells with hyphal coils (Figure 1A), the symbiotic efficiency between *B. striata* and *Tulasnella* was evaluated by comparing the number of symbiotic cells within a protocorm (Yamamoto et al., 2017; Fuji et al., 2020). A two-week-old protocorm contained approximately 81.0±4.0 symbiotic cells with hyphal coils in the control treatment (Figure 1B). The number of symbiotic cells in 1 µM GA treatment was significantly lower (5.4±3.0) than in the control (Figure 1B), while no differences were detected in terms of fungal entry at the suspensor end between the treatments (Supplemental Figure S1B). On the contrary, a GA biosynthesis inhibitor uniconazole-P treatment showed a significant increase in symbiotic cells (135.8±38.2, Figure 1B). Uniconazole-P exhibited a positive effect on hyphal growth, but it was not significant compared to GA treatment (Supplemental Figure S1A). These results resembled those of earlier studies on AM symbiosis in which exogenous GA reduced hyphal colonization and arbuscule formation (El Ghachtouli et al., 1996; Yu et al., 2014; Takeda et al., 2015). The exogenous GA also significantly inhibited seed germination of *B. striata* inoculated with *Tulasnella*. (Figure 1C), in line with previous reports (Chen et al., 2020).

**Figure 1.**
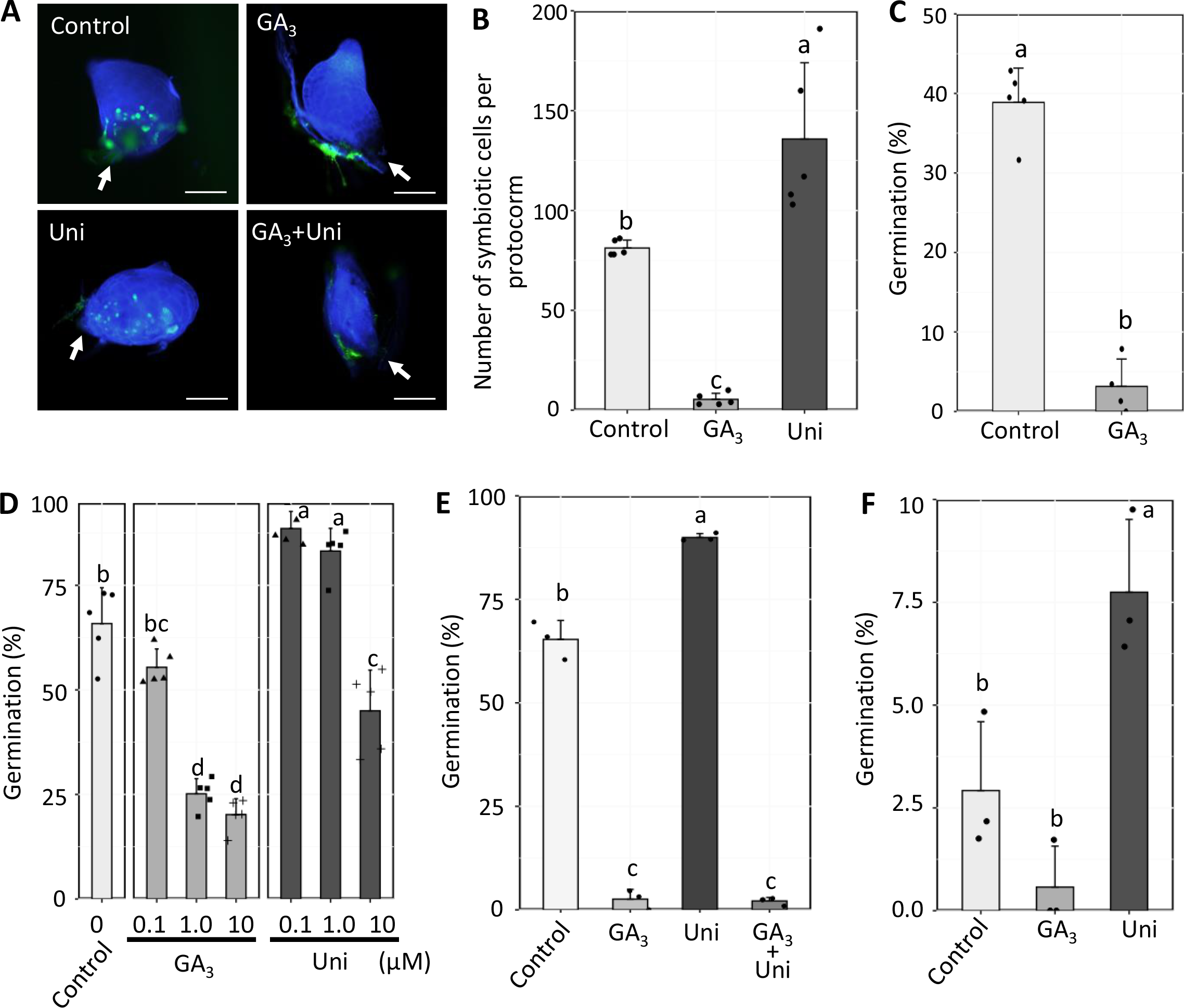
Effects of GA on symbiotically or asymbiotically germinated protocorms. **A)** Symbiotically germinated *B. striata* seeds at two weeks after seeding (WAS) on oatmeal agar (OMA) medium inoculated with *Tulasnella* sp. HR1-1. The images show fungal-infected protocorms stained with calcofluor white (blue) and wheat germ agglutinin-Alexa fluor-488 (green) to visualize the plant cell and fungal structures, respectively. Green fluorescent dots at the suspensor side indicate fungal pelotons. Arrows indicate the suspensor end. Scale bars, 200 µm. **B** and **C)** The number of symbiotic cells **(B)** and germination percentages **(C)** at 2WAS on OMA medium inoculated with *Tulasnella* sp. HR1-1. Symbiotic cells and germinated seeds treated with 0.01% (v/v) ethanol for the control, 1 µM GA3, or 1 µM uniconazole-P (Uni) were counted. Different letters indicate significant differences among treatments on the basis of Tukey’s honest significant difference test (n=5 individual experiments, each containing 10 protocorms, *P* < 0.05) **(B)**. Seed germination was defined as the emergence of a shoot, according to Yamamoto et al. (2017). *P* value was calculated on the basis of Student’s *t*-test (n=4–5 replicate plates, each containing 185±64 seeds) **(C)**. **D** to **F)** Asymbiotically germinated seeds on Hyponex agar medium **(D** and **E)** or filter papers **(F)**. Germinated seeds treated with 0.01% (v/v) ethanol for the control, GA3, or Uni at the indicated concentrations were counted at 2 (d and e) or 3 **(F)** WAS. In e and f, seeds were treated with 1 µM GA3, 1 µM Uni, or 1 µM each of GA3 and Uni. Different characters indicate statistically significant differences on the basis of Tukey’s honest significant difference test (n=3–5 replicate plates, each containing 117±54 seeds, *P* < 0.05). Each bar represents the mean value ± standard deviation. All experiments were independently repeated at least three times with similar results.

Our findings support previous studies that OM symbiosis may share at least some common properties with AM symbiosis (Suetsugu et al., 2017; Miura et al., 2018), but do not support the general notion that GA stimulates seed germination. The *B. striata* seeds were then sown asymbiotically on HA medium with or without GA. The results of germination rates demonstrated that 1 and 10 µM GA treatment, as well as 1 µM ABA, significantly suppressed (25.2±3.6% and 20.2±3.8% in 1 and 10 µM GA, respectively) germination as compared to a control experiment (65.9±8.6%), while 0.1 µM GA showed no significant effect (55.4±4.4%) (Figure 1D and Supplemental Figure S1C). The treatment of 0.1 and 1 µM uniconazole-P showed a promotion effect on seed germination (88.6±4.1% and 83.2±5.4% in 0.1 and 1 µM uniconazole-P, respectively), while 10 µM of uniconazole-P negatively affected the germination (45.0%±9.7%) (Figure 1D). Because uniconazole-P has also been reported to inhibit the biosynthesis of other phytohormones such as ABA and brassinosteroids (Iwasaki and Shibaok, 1991; Kitahata et al., 2005; Saito et al., 2006), a simultaneous treatment test with GA and uniconazole-P was conducted. Asymbiotically germinated seeds that were simultaneously treated with 1 µM GA and 1 µM uniconazole-P showed a lower germination rate than the control as well as with only GA treatment (Figure 1E). Moreover, the germination occurred slightly but significantly even when *B. striata* seeds were grown on filter paper saturated with 1 µM uniconazole-P solution without any nutrition (Figure 1F). Such negative and positive effects of GA and uniconazole-P were not observed in *O. sativa* and *L. japonicus* (Supplemental Figure S2).

To confirm the negative effects of GA on seed germination of other orchids, an asymbiotic germination experiment was conducted using seeds of the other four orchid species: three are terrestrial (*Spiranthes australis*, *Goodyera crassifolia*, and *Cremastra appendiculata* var. *variabilis*), and one is epiphytic (*Vanda falcata* cv. *Beniohgi*). Similar to the results for *B. striata*, exogenous GA treatment decreased the germination percentage of these orchids (Table 1). In addition, commercial plant growth regulators containing substances that inhibit GA biosynthesis showed activity as seed germination stimulators (Supplemental Figure S1D). These results revealed that exogenous GA negatively affected orchid seed germination, indicating that seed germination and fungal colonization are promoted through the inhibition of *de novo* GA biosynthesis. This is different from the seeds with fully differentiated embryos of most photosynthetic plants whose germination is stimulated by GA.

**Table 1.** The effect of gibberellin on orchid seed germination

|  | Period | Control | GA | Uniconazole-P |
| --- | --- | --- | --- | --- |
| *Spiranthes australis* | 28 days | 4.90±1.36<sup>a</sup> | 1.09±0.49<sup>b</sup> | 5.64±0.57<sup>a</sup> |
| *Goodyera crassifolia* | 28 days | 8.51±4.48<sup>a</sup> | 3.79±1.52<sup>a</sup> | 18.34±11.72<sup>a</sup> |
| *Cremastra appendiculata* var. *variabilis* | 60 days | 5.04±1.26 | Not detected | 7.22±4.50 |
| *Vanda falcata* cv. *Beniohgi* | 28 days | 51.30±4.85<sup>a</sup> | 29.89±4.00<sup>b</sup> | Not tested |

<sup>a,b</sup>Different characters indicate statistically significant differences on the basis of the Bonferroni-adjusted pairwise *t*-test (n=3–5, P < 0.05).

### A majority of differentially expressed genes were shared between asymbiotically-and symbiotically germinated protocorms

The negative effects of GA on both seed germination and fungal colonization in *B. striata* led us to hypothesize that orchid germination mechanisms are linked with the symbiotic system. To gain insight into the similarity and differences between asymbiotic and symbiotic germination mechanisms at a molecular level, we performed a time-course transcriptome profiling on asymbiotically or symbiotically germinated protocorms (hereafter APs or SPs, respectively) (Supplemental Table S2). Our previous study showed that *B. striata* grew almost synchronously for the first three weeks with both the germination methods, asymbiotic germination on the HA medium and symbiotic germination with *Tulasnella* on the OMA medium (Yamamoto et al., 2017). In addition, symbiotic protocorms were dominated by fungal pelotons at the early (initiation of hyphal coiling), middle (well-developed hyphal coils), and late (hyphal degradation) stages within one, two, and three weeks after seeding, respectively (Yamamoto et al., 2017; Miura et al., 2018). Up- and down-regulated DEGs were identified from comparisons of each week-old (weeks 1, 2, and 3) protocorms versus seeds at the start of the experiment (week 0) with a |Log2 -FC| threshold ≥ 1 and a false discovery rate threshold < 0.05 (Supplemental Table S3). A total of 10,297 and 14,284 genes in week 1, 10,704 and 15,606 genes in week 2, and 10,835 and 15,244 genes in week 3 had a significantly higher expression compared with week 0 in APs and SPs, respectively. Among these DEGs, 8,663, 8693, 8217 genes were common in both APs and SPs comparison sets, which covered 38.6%–52.5% of the total number of DEGs from weeks 1–3 (Figure 2A). A total of 6,199 and 5,998 genes in week 1, 6,136, and 6,092 genes in week 2, and 5,858 and 6,020 genes in week 3 had significantly lower expression than week 0 in APs and SPs, respectively (Figure 2A). Among these, 4,783, 4,410, and 0 genes were common in weeks 1, 2, and 3, respectively (Figure 2A).

**Figure 2.**
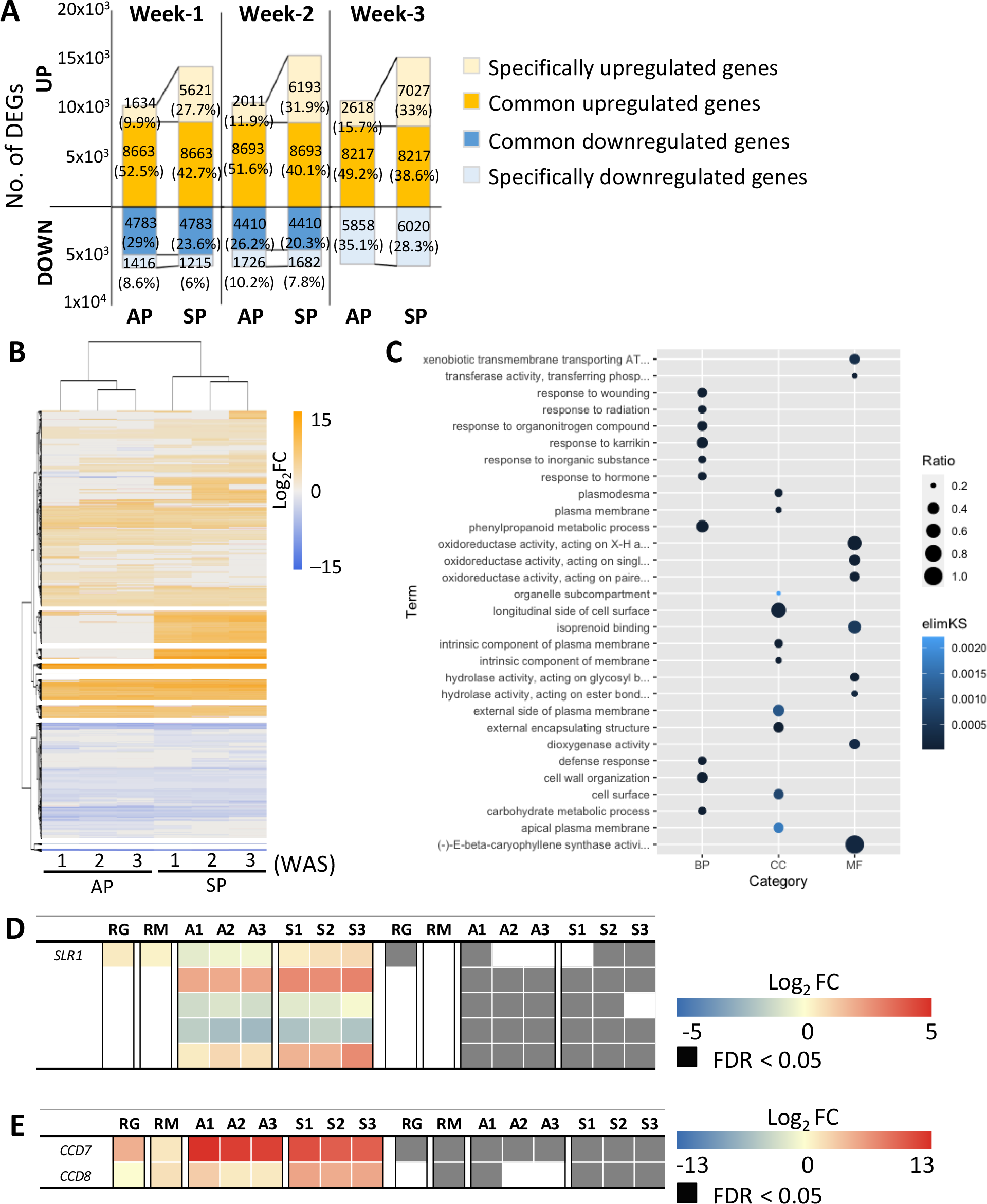
Transcriptome analysis of asymbiotically and symbiotically germinated *Bletilla striata*. **A)** The bar chart of the number of differentially expressed genes (DEGs). Gene expression levels of asymbiotically germinated protocorms (AP) and symbiotically germinated protocorms (SP) at 1–3 weeks after seeding were compared to week-0 seeds. **B)** Hierarchical clustering of DEGs. The method of k-means clustering was used to identify similarities in expression patterns among AP and SP. The heatmap was drawn by the pheatmap package in R. **C)** Gene-ontology (GO) enrichment analysis of shared overexpressed genes between AP and SP at week 1. The most significant ten terms of each category, biological process (BP), cellular component (CC) and molecular function (MF), are shown on the basis of the elim-Kolmogorov-Smirnov method in the topGO package in R. All significant terms were presented in Table **S4**. **D** and **E)** The expression patterns of *SLENDER RICE1* (*SLR1*) orthologs containing DELLA domain **(D)** and carotenoid cleavage dioxygenase genes (*CCD7* and *CCD8*) involved in strigolactone biosynthesis **(E)** from RNA-seq data. The blue–red heatmap on the left shows the expression patterns of the selected genes on the basis of Log2-fold change (FC). Log2FC was calculated between time points; 0-week seeds versus 1–3-week protocorms (A1–A3 and S1–S3). “A” and “S” represent asymbiotic and symbiotic germination, respectively. “RG” and “RM” indicates germinated seeds and arbuscular mycorrhizal roots of rice, respectively. The right panel displays false discovery rates (FDR).

To compare the transcription profiles of seed germination and mycorrhizal symbiosis in other plant species, DEGs were identified using the previous transcriptome data of seed germination on day 0 versus day 2 and AM roots versus non-AM roots of *O. sativa* (Narsai et al., 2017; Kobae et al., 2018). The rice transcriptome data showed that shared DEGs between seed germination and AM symbiosis covered 0.7%–23.9% of the total DEGs (Supplemental Figure S3A).

Hierarchical clustering based on the *B. striata* expression profiles showed that the transcripts were divided into AP and SP, as expected (Figure 2B). Transcripts at week 1 exhibited different expression patterns from those of weeks 2 and 3 (Figure 2B). Additionally, over 50% of DEGs were shared between AP and SP at week 1 (Figure 2A), and GA restricted peloton formation but still allowed the fungal entry at the suspensor end (Supplemental Figure S1A). On the basis of these results, it was expected that the stage of forming a shoot apex in APs or fungal colonization following fungal entry from the suspensor end to cortical cells in SPs is the key to understanding mycoheterotrophic germination. Then, the shared and specific DEGs at week 1 were annotated into three categories (“Biological process,” “Cellular component,” and “Molecular function”) after GO enrichment analysis with a cut-off *P*-value 0.01 (Supplemental Table S4). The overexpressed genes in the shared DEGs were classified into 95 functional GO terms, including response to (in)organic substances such as “response to organonitrogen compound (GO:0010243)”, “response to karrikin (GO:0080167)”, “response to inorganic substance (GO:0010035)”, and “response to hormone (GO:0009725)”, and metabolism of secondary metabolites, such as “phenylpropanoid metabolic process (GO:0009698)”, “(-)-E-beta-caryophyllene synthase activity (GO:0080016)” (Figure 2C).

Specifically overexpressed genes in APs and SPs were assigned to 14 and 82 terms, respectively (Supplemental Table S4). The differently overrepresented GO terms related to nitrogen response and metabolism (“cellular nitrogen compound metabolic process (GO:0034641)” in APs and “response to organonitrogen compound (GO:0010243)” in SPs) (Supplemental Figure S3B and C) were consistent with the differences in applied nitrogen form between APs as inorganic matter in the culture medium and SPs as organic matter from the fungal partner. In addition, different GO terms related to secondary metabolism were overrepresented in AP (“cellular aromatic compound metabolic process [GO:0006725]”) and SP (“cellular alcohol metabolic process [GO:0044107]”) (Supplemental Figure S3B and C). In the SP, GO terms associated with the transporter activity, such as “intracellular transport (GO:0046907)” and “efflux transmembrane transporter activity (GO:0015562),” were specifically overrepresented, probably reflecting nutrient transport between plants and fungi (Supplemental Figure S3C).

To understand the intrinsic metabolic and signaling of GA during seed germination and fungal colonization, we performed the Kyoto Encyclopedia of Genes and Genomes analysis on the common DEGs, termed as “response to hormone (GO:0009725)”. This analysis revealed that the ent-kaurene oxidase gene (TRINITY_DN56117_c0_g1_i2_g.199526_m.199526), which is involved in the GA synthesis, and the *DELLA* gene (TRINITY_DN29017_c0_g1_i1_g.90488_m.90488), which is a negative regulator of GA signaling, were detected as commonly upregulated genes in APs and SPs (Supplemental Fig. S4). This result was inconsistent with *Arabidopsis* seed germination, in which *DELLA* genes were either constantly expressed or gradually decreased from 12 h to 2 days after imbibition (Tyler et al., 2004), and the rice *DELLA* ortholog *SLENDER RICE1* expression was almost not altered during seed germination (Log2FC of 0.49) (Figure 2D). In cereal seeds, the α-amylase gene acts as one of the downstream genes under GA-mediated seed germination (Gubler et al., 2002). A GA-regulated transcriptional factor, GAMYB, activated α-amylase gene expression and was promoted by GA-triggered degradation of DELLA protein *SLENDER1* in the barley aleurone (Gubler et al., 2002). Consistently, the expression of α-amylase genes was markedly increased in germinated rice seeds (Supplemental Figure S3D). In contrast, there were no significant differences between APs at week 1 and week 0 seeds (Supplemental Figure S3D). In addition, according to the other transcriptomic studies of symbiotically germinated *D. officinale* inoculated with *Tulasnella* sp. Pv-PC-2-1 (Wang et al., 2018), common upregulated DEGs at week 1 included two carotenoid cleavage dioxygenases (*CCD7* and *CCD8*) genes, which are necessary for strigolactone (SL) biosynthesis (Figure 2E). Previous studies reported that the exogenous GA inhibited SL biosynthesis and exudation in *O. sativa*, *L. japonicus*, and *Eustoma grandiflorum* roots (Ito et al., 2017; Tominaga et al., 2021). Consistently, both *CCD7* and *CCD8* were upregulated in AM-colonized rice roots (Log2FC of 1.43 and 1.89, respectively), whereas only *CCD7* was detected as significantly overexpressed in germinated seeds in rice (Figure 2E). These results indicate that although GA biosynthesis is activated during seed germination in *B. striata* as well as other plant species, such as *Arabidopsis*, rice, and barley (Kaneko et al., 2002; Gubler et al., 2008; Dekkers et al., 2013; Urbanova and Leubner-Metzger, 2018), simultaneously, other factors, such as DELLA proteins, could inhibit GA signaling during *B. striata* seed germination.

### Bioactive GA is converted to the inactive form during *B. striata* seed germination

Given that GA negatively controls both seed germination and fungal colonization, and the APs and SPs shared a large number of DEGs, including GA biosynthesis and signaling genes, it is expected that *B. striata* optimize GA production during seed germination to coordinate both seed germination and fungal colonization. The transcriptome analysis showed that GA 2-oxidase genes (*GA2oxs*) encoding enzymes that convert bioactive GAs to inactive forms and GA 3- and 20-oxidase genes (*GA3oxs* and *GA20oxs,* respectively) encoding the biosynthesis enzymes to produce the active form of GAs were significantly upregulated in APs and SPs (Figure 3A). These significant changes in the expression of *GA2oxs* and *GA20ox* genes were also detected in rice seed germination, but moderately found in AM-colonized rice roots (Figure 3A). To reveal more details of the regulation of GA metabolism, gene expression analysis was performed by quantitative RT-PCR. In APs, the expression of a *BsGA2ox* was 19 and 45 times upregulated at week 1 and week 2, respectively, compared to week 0, which was the start of the experiment, while week 1 was not significant in this experiment (Figure 3B). Similarly, the expression of *BsGA3ox* was 12 and 40 times higher at weeks 1 and 2, respectively, whereas *BsGA20ox* was not significantly altered. In SPs, the expression of *BsGA2ox*, *BsGA3ox*, and *BsGA20ox* was significantly upregulated, especially *BsGA2ox* expression was dramatically increased (116 and 113 times higher at weeks 1 and 2, respectively) after seeding (Figure 3C). These results provide an insight into the mechanism of *B. striata* seed germination and fungal colonization: The GA metabolism is stimulated, especially the bioactive GA is actively converted to the inactive form.

**Figure 3.**
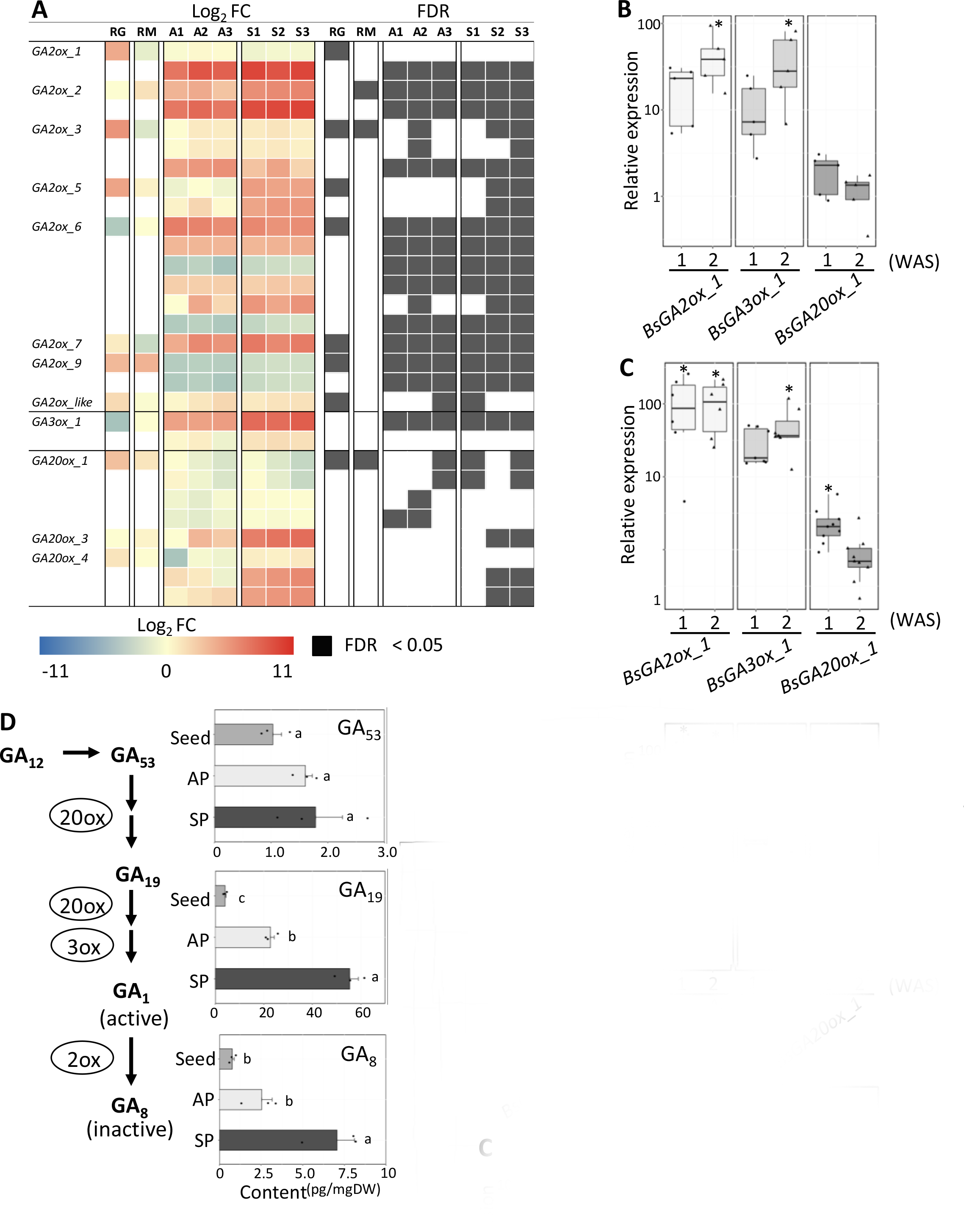
Expression of gibberellin biosynthesis and metabolism genes during asymbiotic and symbiotic germination. **A)** The expression patterns of GA biosynthesis and metabolism genes from RNA-seq data. The blue–red heatmap on the left shows the expression patterns of the selected GA-related genes on the basis of Log2-fold change (FC). Log2FC was calculated between time points; 0-week seeds versus 1–3-week protocorms (A1–A3 and S1–S3). “A” and “S” represent asymbiotic and symbiotic germination, respectively. “RG” and “RM” indicates germinated seeds and arbuscular mycorrhizal roots of rice, respectively. The right panel displays false discovery rates (FDR). **B** and **C)** Quantitative RT-PCR of gibberellin biosynthesis and metabolism genes. Relative expression analysis was performed with total RNA isolated from asymbiotically **(B)** or symbiotically **(C)** germinated *B. striata* at different time points (week 0 seeds to protocorms at two weeks after seeding (WAS)). The relative expression values were determined using the relative standard curve method. The fold change in expression is relative to 0-week-old seeds (expression level = 1). Asterisks indicate significant differences compared to the 0-week-old seeds using a Bonferroni-adjusted pairwise *t*-test (n=5–9, * *P* < 0.05). **D)** The content of endogenous gibberellins (GAs) during seed germination. The GAs were detected in seeds at week 0 (Seed), asymbiotically and symbiotically germinated protocorms at week 2 (AP and SP, respectively). Different characters indicate statistically significant differences on the basis of the Bonferroni-adjusted pairwise *t*-test (n=3, *P* < 0.05). Each bar represents the mean value ± standard error.

To further confirm these findings, we conducted a quantitative analysis of endogenous GA in APs and SPs at week 2 and week 0 seeds using LC-MS/MS. The study showed endogenous bioactive GA precursors (GA19 and GA53) and an inactivated GA (GA8) in the 13-hydroxylation pathway (Figure 3D), while bioactive GA (GA1) could not be detected due to the detection limit of the instrument. In APs, the amount of GA19 was significantly increased (5.6 times), and the GA8 was 3.2 times higher but not significant compared with that in seeds. The GA53 was slightly higher (1.5 times) in APs than in seeds, but not significant. The quantification analysis also showed that in SPs, the amount of GA8 and GA19 was significantly increased compared to the seeds, whereas no significant difference was observed in the amount of GA53 accumulated during the experimental period. The gene expression and GA quantification results revealed that the biosynthetic pathway from the GA precursor GA19 to the inactive form GA8 was activated during both seed germination and fungal colonization, especially, BsGA2ox strongly expressed in SPs actively converted bioactive GAs to inactive forms during symbiotic germination.

### Asymbiotic germination induced the expression of orchid mycorrhizal marker genes

Our transcriptome results of shared gene expression patterns between APs and SPs were reminiscent of those showing that the artificial media for asymbiotic germination would mimic something in the field probably provided by fungi (Rasmussen, 1992; Eriksson and Kainulainen, 2011). In other words, these findings have led to the hypothesis that symbiotic machinery is activated automatically during seed germination even without fungi. To determine whether the symbiosis-signaling pathway is activated during asymbiotic germination, the expression levels of rice AM colonization marker homologs (Gutjahr et al., 2008), part of which were identified as OM markers in *B. striata* (Miura et al., 2018), and the mycorrhiza-specific phosphate transporter *PT11* gene were compared between rice and *B. striata*. The previous rice transcriptome data showed nine significantly highly expressed marker genes in AM-forming roots, and only one in germinated seeds (Figure 4A). Our time-course transcriptome analysis revealed that 11 AM marker homologs were significantly highly expressed in SPs (Figure 4A). Similarly, the high expression of nine AM marker genes was observed in APs (Figure 4A). The *PT11* expression was significantly induced in both APs and SPs as well as AM-forming roots, but not in rice-germinated seeds (Figure 4A). Consistently, quantitative RT-PCR analysis of AP showed that the expression of six of eight OM marker genes (*BsAM1*, *BsAM2*, *BsAM11*, *BsAM14*, *BsAM20*, *BsAM34*) was upregulated during germination despite the absence of fungus, whereas *BsAM34* was not significant (Figure 4B).

**Figure 4.**
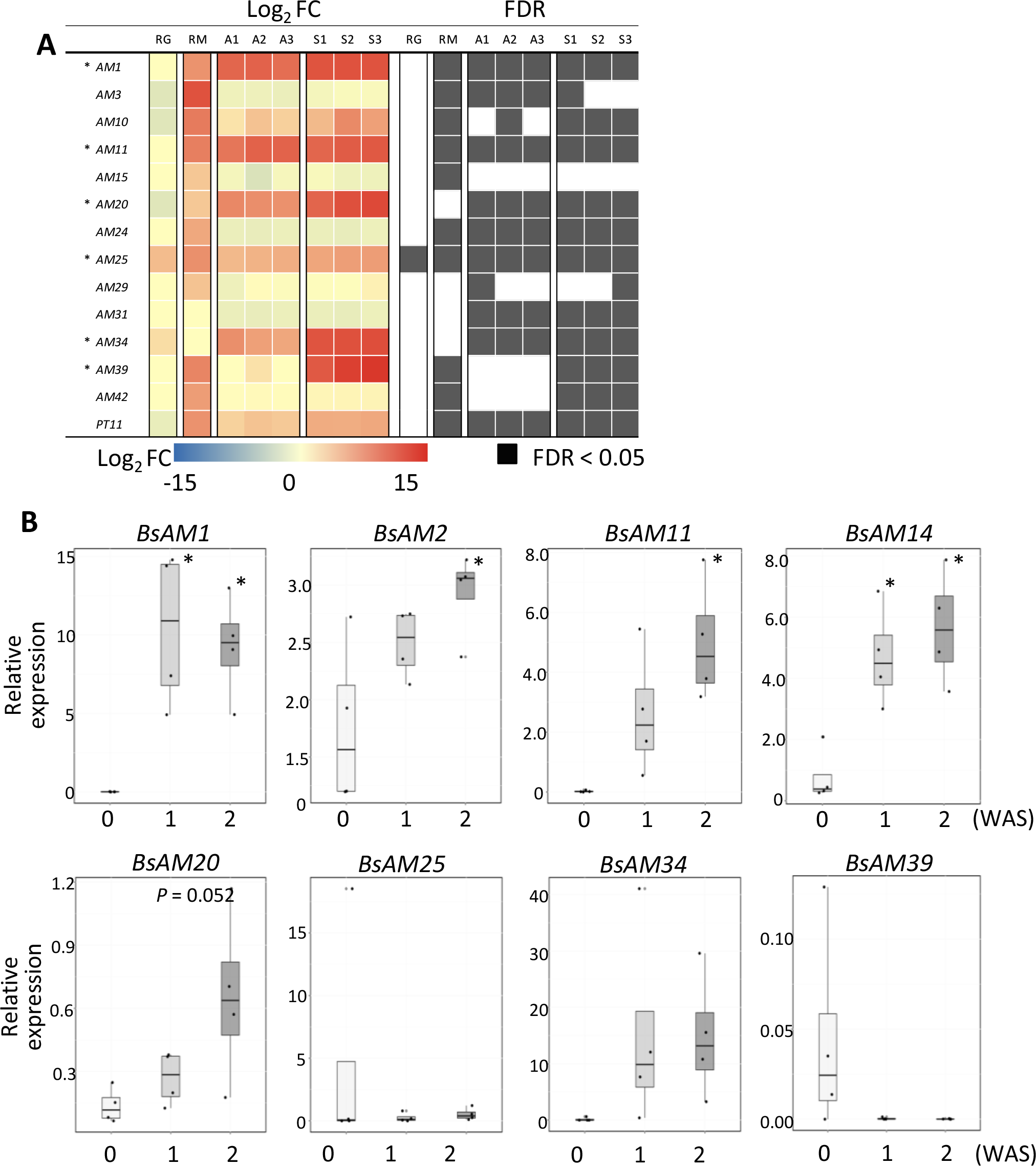
Expression of symbiosis marker genes during asymbiotic germination. **A)** The expression patterns of symbiosis marker genes of rice and *Bletilla striata* from RNA-seq data. The heatmap on the left indicates the expression patterns of the arbuscular mycorrhizal symbiosis marker genes (Gutjahr et al., 2008) and the mycorrhiza-specific phosphate transporter *PT11* gene on the basis of Log2-fold change (FC). In rice, Log2FC was calculated between day 0 seeds and germinated seeds at two days after seeding (RG) and between non-colonized roots and arbuscular mycorrhizal (AM) roots (RM). In *B. striata*, Log2FC was computed for week 0 seeds versus 1– 3-week protocorms (A1–A3 and S1–S3). “A” and “S” represent asymbiotic germination and symbiotic germination, respectively. The right panel displays false discovery rates (FDR). Asterisks indicate the orchid mycorrhizal (OM) symbiosis marker genes (Miura et al., 2018). **B)** Quantitative RT-PCR of OM symbiosis marker genes. Relative expression analysis was performed with total RNA isolated at different time points (week 0 seeds to germinated protocorms at two weeks after seeding (WAS)). The relative expression values were determined using the relative standard curve method. Asterisks indicate significant differences compared to the week-0-old seeds using the Bonferroni-adjusted pairwise *t*-test (n=4, * *P* < 0.05).

## Discussion

It is generally accepted that seed dormancy and germination are determined by the interactive effects between different phytohormones such as ABA and GA, and that GA is necessary for seed germination (Miransari and Smith, 2014). However, our results show that exogenous GA, as well as ABA, significantly inhibits the germination of *B. striata* seeds with undifferentiated embryos, even on a nutrition-free medium. These imply the existence of a specific mechanism of GA signaling in *B. striata* seed germination (Figure 5). This is consistent with some earlier studies of terrestrial and epiphytic orchids, which reported that GA had a negative or no effect during germination (Van Waes and Debergh, 1986; Miyoshi and Mii, 1995; Chen et al., 2020). These results indicate that orchid species broadly conserve this unique GA signaling pathway.

**Figure 5.**
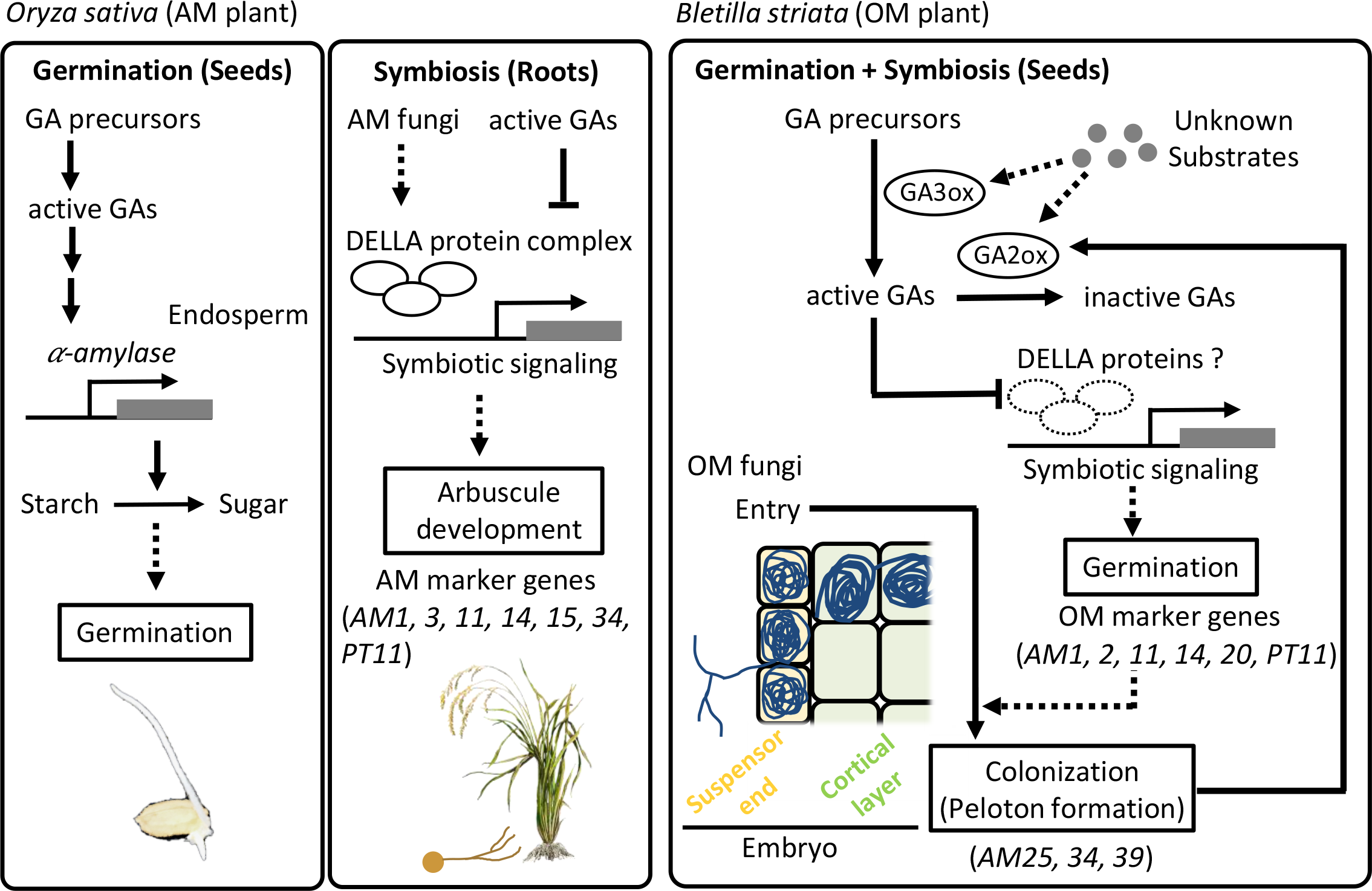
Proposed model for seed germination mechanism in orchids. Gibberellin (GA) stimulate seed germination of arbuscular mycorrhizal (AM) plants, such as *Oryza sativa*, by inducing the expression of α-amylases necessary for the utilization of starch stored in the endosperm (Kaneko et al., 2002). After root development, the mutual relationships between plants and AM fungi are established in the roots. Exogenous-treated GA inhibits fungal colonization in rice through the degradation of DELLA proteins. In *Bletilla striata*, an orchid mycorrhizal plant, exogenous treated GA inhibits seed germination and fungal colonization via unknown mechanisms. When seed germination occurs, environmental factors probably derived from OM fungi stimulate the expression of GA metabolic genes such as GA3ox and GA2ox, leading to symbiotic signaling even without fungi. Orchid mycorrhizal fungi form pelotons in the cortical layer, which promotes *GA2ox* gene expression.

Connecting multiple signaling pathways has the potential to acquire new adaptive functions (True and Carroll, 2002). Previous studies on model plants have reported that the gene regulatory mechanisms stimulated by circadian and environmental cues contribute to the resistance against a plant pathogen (Wang et al., 2011) and the facilitation of AM symbiosis (Umehara et al., 2008; Balzergue et al., 2011; Kretzschmar et al., 2012). These studies have established the concept of molecular links between intrinsic or extrinsic cues as an input and signaling pathways that seem far apart from these cues as an output. Our results showed that GA has lost its role as an activator of α-amylase gene expression during orchid seed germination. Instead, an exogenous germination stimulator probably derived from OM fungi, but not the fungal colonization itself activated symbiosis-related genes shared with AM symbiosis through GA inactivation, leading to simultaneous control of seed germination and fungal colonization. In the AM symbiosis signaling mutant *ccamk-1* of rice wherein the mutation blocked AM colonization at the root epidermis, no increase in AM marker expression except for *OsAM1* and *OsAM2*, was observed in AM-inoculated roots (Gutjahr et al., 2008). The enhancement of the expression of *B. striata* OM marker genes in APs could result from the activation of symbiosis signaling by germination stimulators, not by fungal colonization. These results suggest that orchids auto-activate the mycorrhizal symbiosis pathway through GA inactivation during seed germination to accept the fungal partner immediately. In addition, the results showing that the number of DEGs identified from SP transcripts was higher than that from AP, one of the OM marker genes *BsAM39* was expressed only in SP, and the *GA2ox* expression level was more stimulated in SPs than in AP suggest that the OM symbiosis pathway is activated in each of the two steps of symbiotic germination: the initiation of seed germination and fungal colonization.

In contrast to the exogenous GA effect, the GA-biosynthesis inhibitor uniconazole-P positively regulates *B. striata* seed germination, although the germination rate decreased at high concentrations, probably because uniconazole-P affects not only GA biosynthesis but also the metabolism of some phytohormones (Izumi et al., 1988; Iwasaki and Shibaok, 1991). Previously, it was thought that orchid seeds need to be colonized by fungi to germinate even *in vitro* (Arditti, 1967). However, since Knudson (1922) formulated the asymbiotic germination method, the fact that germination occurs when seeds are placed in a relevant medium containing appropriate substances, such as sugars, mineral salts, vitamins, amino acids, and phytohormones without fungus is broadly accepted (Arditti, 1967). Thus, artificial media for asymbiotic germination would mimic something in the field probably provided by fungi (Rasmussen, 1992; Eriksson and Kainulainen, 2011). Our results support the hypothesis that orchid germination is stimulated by environmental factors instead of fungal infection because exogenously treated uniconazole-P stimulated *B. striata* germination even without any other nutrients and fungi. Moreover, the uniconazole-P acts as a germination stimulator commercially accessible and usable by gardeners, offering fundamental knowledge to the development of conservation and restoration methods of orchids facing extinction (Swarts and Dixon, 2009).

Although the actual influencing environmental factor(s) are still unknown, our results suggest that the unknown factors induce the GA metabolic pathway, resulting in the stimulation of seed germination and fungal colonization. In both APs and SPs, the expression of *GA2ox* was significantly increased. Consistently, the inactivated GA, GA8, accumulates in APs and SPs but is not statistically different between APs and seeds before seeding. A previous transcriptomic study has also reported high expression of genes related to the GA-GID1-DELLA signaling module, including *GA2ox* and *GA20ox*, in the protocorms of *Anoectochilus roxburghii* inoculated with unknown fungal species (Liu et al., 2015). These results indicate that the bioactive GA is actively converted to the inactive form during both seed germination and fungal colonization in orchids. The negative effect of bioactive GA in symbiotic germination resembles the results of earlier studies on AM symbiosis, which reported that GA signaling negatively affects AM fungal colonization and development (Floss et al., 2013; Foo et al., 2013; Takeda et al., 2015). Our previous study found that the key components of AM symbiosis and their concomitant expression pattern are shared in OM symbiosis (Miura et al., 2018). Given that the fungal partner(s) is necessary for orchid seed germination under natural conditions, these findings suggest that the GA is converted from the active form to the inactive form to establish and maintain symbiotic associations during seed germination.

In orchids, the endosperm was reduced, and the undifferentiated embryo evolved (Eriksson and Kainulainen, 2011). Orchid seeds containing almost no or few resources are tiny and require the presence of compatible fungi to obtain nutrients, resulting in mycoheterotrophy (Eriksson and Kainulainen, 2011). The evolutionary process of dust-like seeds in such mycoheterotrophic plants is supposed to be complex because many drivers behind fungal specificity, chemical interaction, metabolite transport, and plastid genome evolution are responsible for this unique plant—fungus relationship (Eriksson and Kainulainen, 2011; Merckx, 2013). One of the potential conditions under which initial mycoheterotrophy has evolved could be associated with the acquisition of the ability to utilize exogenous nutrients during seed germination. Our findings imply that orchids have co-opted the AM symbiotic signaling pathway for GA-mediated germination to coordinate these two simultaneous events. These molecular links between plant development and symbiosis signaling will provide clues to the mechanisms of the evolutionary shift of the autotrophic ancestor to the mycoheterotrophic orchids.

## Acknowledgments

We thank Mr. Ikuo Nishiguchi and the National BioResource Project (Legume Base) for providing *V. falcata* and *L. japonicus* seeds, respectively. We also thank the Data Integration and Analysis Facility, National Institute for Basic Biology (NIBB) for supporting the RNA sequencing and providing computational resources. The illustrations were modified and/ created with images from TogoTV (©2016 DBCLS TogoTV / CC-BY-4.0). This work was supported by the NIBB Cooperative Research Programs (Next-Generation DNA Sequencing Initiative: 15-825, 16-430; 17-430, 18-441, 19-433, 20-407, 21-301, 22NIBB403), the Japan Society for the Promotion of Science (JSPS) KAKENHI (Grant Number 15K14550, 18J01755), JSPS Research Fellowships for Young Scientists, and the Tottori Prefecture Research Fund for the Promotion of Environmental Academic Research. We thank Enago (www.enago.jp) for the English language review.

## Author contributions

CM and HK conceived and designed the analysis. CM, YF, TY, MH, KS, TY, and MY contributed to sample preparation. CM, YF, TY, YK, MH, and KY performed the experiments and analyzed the sequencing data. KS, TY, MS, SS, and MY helped supervise the project and contributed to the interpretation of the results. CM and HK wrote the paper. HK supervised the project. All authors approved the final manuscript.

## Data availability

Nucleotide sequence data from the RNA-seq analysis in this study have been deposited in the DDBJ BioProject under the accession number PRJDB14881. The authors declare no competing financial interests. Correspondence and requests for materials should be addressed to H.K.

## Supplemental data

**Supplemental Figure S1** The effect of phytohormones on fungal growth and seed germination.

**Supplemental Figure S2** The effect of gibberellin (GA) on seed germination of *Oryza sativa* and *Lotus japonicus*.

**Supplemental Figure S3** Transcriptome analysis of *Oryza sativa* and *Bletilla striata*.

**Supplemental Figure S4** The Kyoto Encyclopedia of Genes and Genomes analysis of shared overexpressed genes between asymbiotically and symbiotically germinated *B. striata* at week-1.

**Supplemental Table S1** List of primers used in this study

**Supplemental Table S2** RNA-sequencing summary of *Bletilla striata*

**Supplemental Table S3** Lists of differentially expressed genes

**Supplemental Table S4** Lists of overrepresented gene ontology terms

