## Supplemental Figure 1-4 for "Orchid seed germination through auto-activation of mycorrhizal symbiosis signaling regulated by gibberellin"

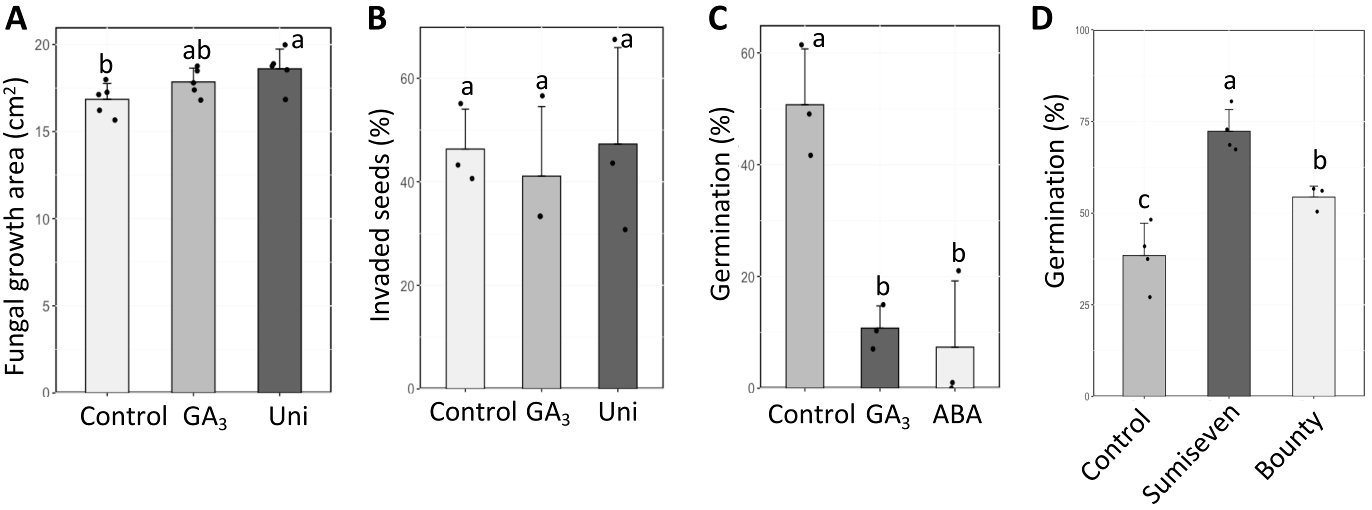


**Supplemental Figure S1** The effect of phytohormones on fungal growth and seed germination. **A)** The effect of gibberellin and its inhibitor on fungal growth. The orchid mycorrhizal fungus *Tulasnella* sp. was inoculated on oatmeal agar (OMA) medium and incubated for seven days. The fungal growth area was measured using the imageJ software. Different characters indicate statistically significant differences on the basis of the Bonferroni-adjusted pairwise *t*-test (n=5, $P < 0.05$). **B)** Fungal entry to the suspensor end of the embryo. *Bletilla striata* seeds inoculated with *Tulasnella* sp. were cultured on (OMA) medium for two days. Fungal-infected seeds were observed under a fluorescence microscope (Leica DM2500, Wetzlar, Germany) after staining with wheat germ agglutinin-Alexa fluor-488. The seed where the fungus enters the suspensor end was determined as an invaded seed. The same letter represents no significant difference on the basis of the Bonferroni-adjusted pairwise *t*-test (n=3, $P < 0.05$). **C)** *B. striata* seeds were germinated on a Hyponex agar medium with gibberellin (GA) and abscisic acid (ABA). Germination rates were measured two weeks after seeding. Each bar represents the mean value ± standard error from three biological repeats. Different characters indicate statistically significant differences on the basis of the Bonferroni-adjusted pairwise *t*-test ($P < 0.05$). Control, Mock (0.01% ethanol)-treated seeds; $GA_3$, 1 μM gibberellin A3; ABA, 1 μM abscisic acid. Each bar represents the mean value ± standard deviation. All experiments were independently repeated two times with similar results. **D)** *B. striata* seeds were germinated on a Hyponex agar medium with plant growth regulators. Germination rates were measured two weeks after seeding. Each bar represents the mean value ± standard deviation according to five biological repeats. Different characters indicate statistically significant differences on the basis of the Bonferroni-adjusted pairwise *t*-test ($P < 0.05$). Control, mock (0.01% ethanol)-treated seeds; Sumiseven, 10 ppb Sumiseven P (Sumitomo Chemical, Tokyo, Japan) containing Uniconazole-P as an active ingredient; Bounty, 10 ppb Bounty Flowable (Syngenta Japan, Tokyo, Japan) containing paclobutrazol.

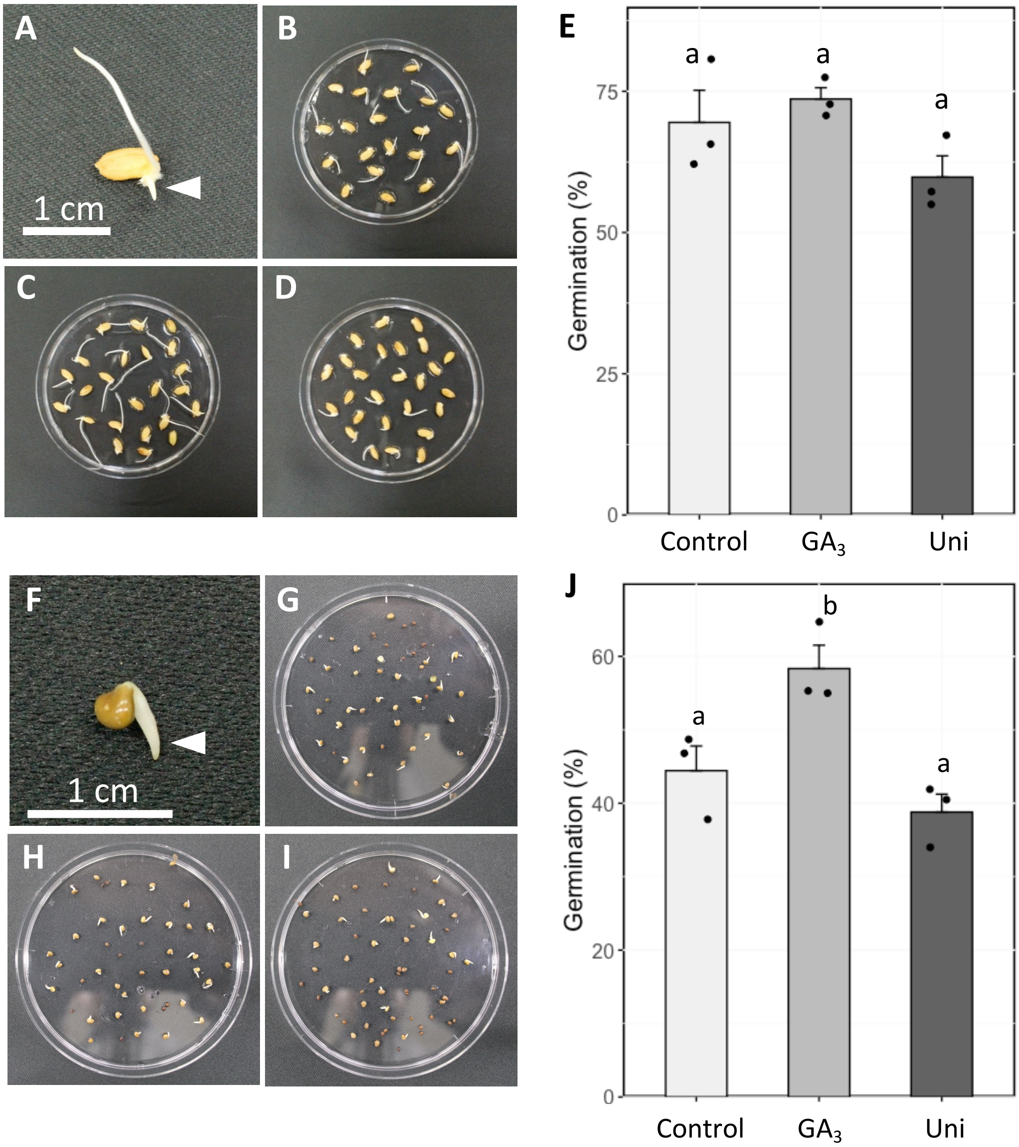


**Supplemental Figure S2** The effect of gibberellin (GA) on seed germination of *Oryza sativa* and *Lotus japonicus*. The seeds of *O. sativa* **(A–E)** and *L. japonicus* **(F–J)** were germinated on a filter paper with $GA_3$ or uniconazole-P. Germination percentages of *O. sativa* **(E)** or *L. japonicus* **(J)** were measured three or two days after seeding, respectively. Seed germination was defined as the emergence of the primary root (arrowhead). Each bar represents the mean value ± standard error from three biological repeats. Different characters indicate statistically significant differences (Tukey-Kramer test, n=3, $P < 0.05$). Control, Mock (0.01% ethanol)-treated seeds **(B, G)**; $GA_3$, 1 μM $GA_3$-treated seeds **(C, H)**; Uni, 1 μM Uniconazole-P-treated seeds **(D, I)**.

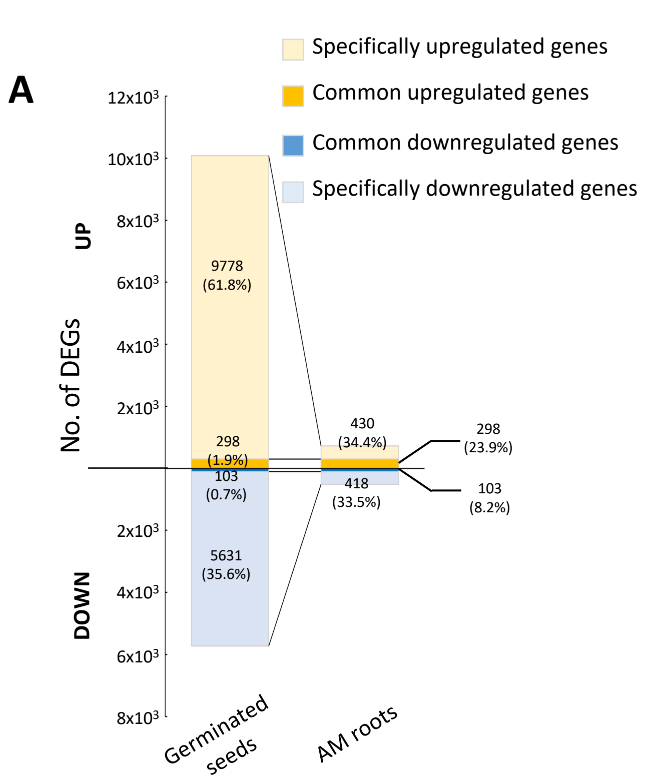


**Supplemental Figure S3** Transcriptome analysis of *Oryza sativa* and *Bletilla striata*. **A)** The bar chart of the number of differentially expressed genes (DEGs) in germinated seeds and arbuscular mycorrhizal (AM) roots of rice. Gene expression levels were compared between 2-day-old germinated seeds and week 0 seeds, and AM-colonized and non-colonized roots. **B** and **C)** Gene ontology (GO) enrichment analysis of specifically overexpressed genes between asymbiotically **(B)** and symbiotically- **(C)** germinated *B. striata* at week 1. The most significant 10 terms of each category, biological process (BP), cellular component (CC), and molecular function (MF), are shown on the basis of the elim-Kolmogorov-Smirnov method in the topGO package in R. All significant terms were presented in Supplemental Table S3. **D)** The expression patterns of α-amylase orthologs from RNA-seq data. The blue–red heatmap on the left shows the expression patterns of the selected genes on the basis of $Log_2$-fold change (FC). $Log_2FC$ was calculated between time points; 0-week seeds versus 1–3-week protocorms (A1–A3 and S1–S3). "A" and "S" represent asymbiotic and symbiotic germination, respectively. "RG" and "RM" indicates germinated seeds and arbuscular mycorrhizal roots of rice, respectively. The right panel displays false discovery rates (FDR).

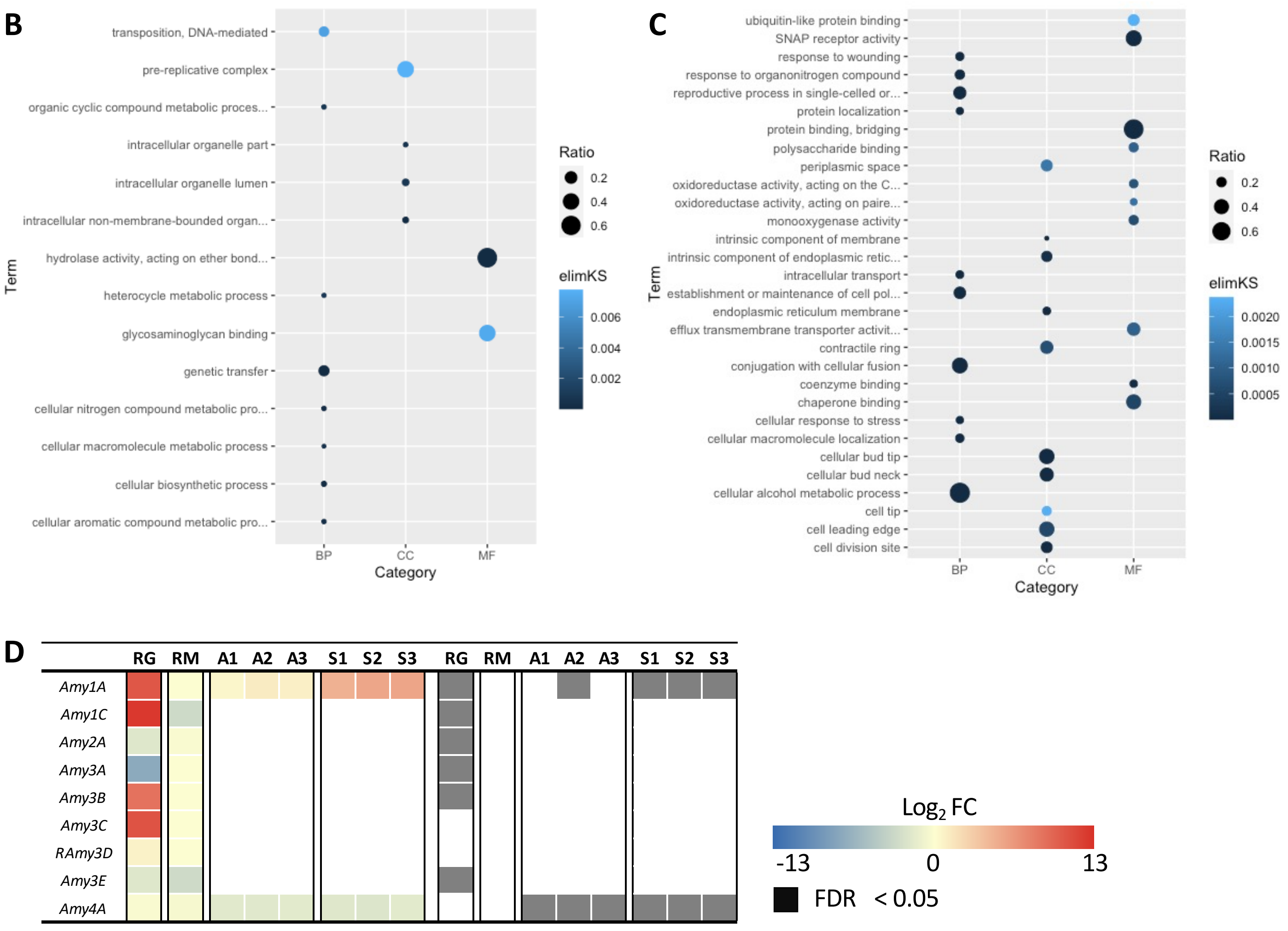

A

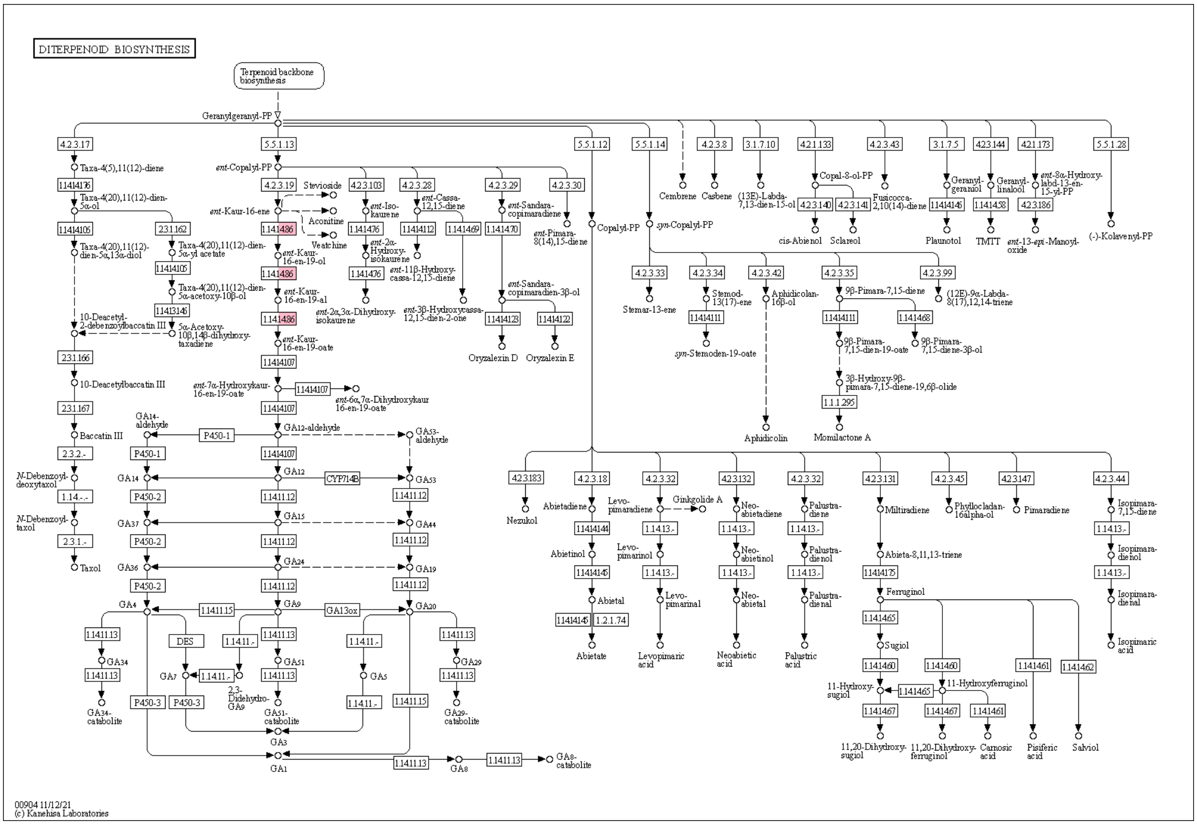


B

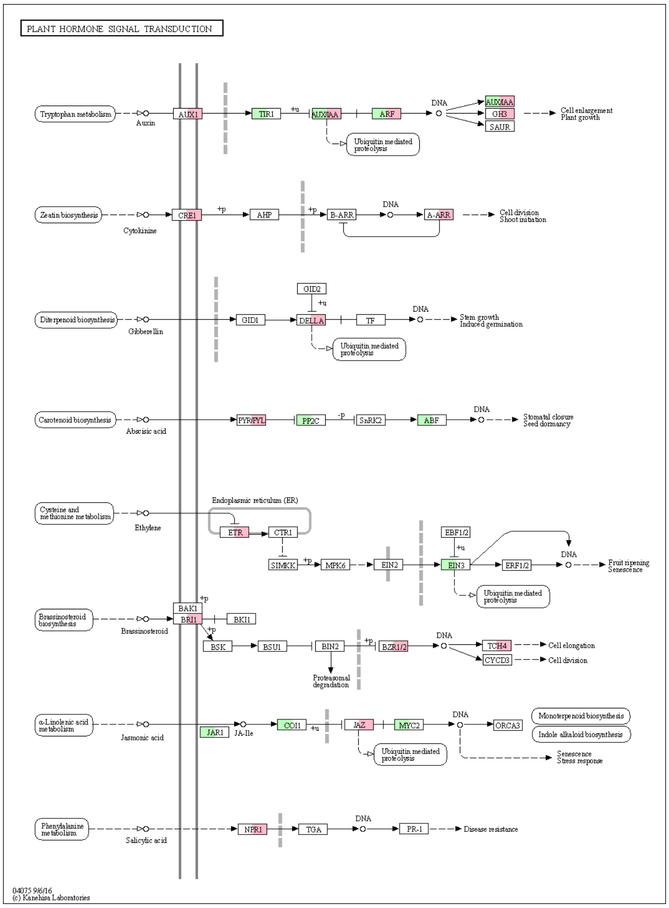


**Supplemental Figure S4** The Kyoto Encyclopedia of Genes and Genomes (KEGG) analysis of shared overexpressed genes between asymbiotically- and symbiotically-germinated *B. striata* at week-1. The KEGG analysis was performed on the common DEGs, especially termed as "response to hormone (GO:0009725)" after annotation with KEGG orthology by using BlastKOALA (https://www.kegg.jp/blastkoala/). The images show the intrinsic metabolic **(A)** and signaling **(B)** of gibberellin. The red color indicates up-regulated genes, and the green indicates down-regulated genes.
